# Road encroachment mediates species occupancy, trait filtering and dissimilarity of songbird communities

**DOI:** 10.1101/2022.01.14.476305

**Authors:** Fernando Ascensão, Marcello D’Amico, Eloy Revilla, Henrique M. Pereira

**Affiliations:** Centre for Ecology, Evolution and Environmental Changes (cE3c), Faculdade de Ciências, Universidade de Lisboa, Lisboa, Portugal; Department of Conservation Biology, Estación Biológica de Doñana (EBD-CSIC), Sevilla, Spain; German Centre for Integrative Biodiversity Research (iDiv) Halle-Jena-Leipzig, Germany

## Abstract

Assessing the road effects on biodiversity is challenging because impacts may depend on both wildlife responses to roads and on the spatial arrangement of roads. We questioned whether an increase in road encroachment i.e., the advancement of roads into non-urban areas, leads to significant changes (positive and negative) in species occurrence, and if so, whether those changes are linked to specific traits related to perturbation sensitiveness, therefore acting as filters that increase the community compositional dissimilarity. Using a large citizen-science dataset of point-counts performed throughout Iberian Peninsula (4,459 unique survey sites), we modelled the effect of road density on the occurrence of common songbirds (n=58), while accounting for potential confounding effects of environment and survey effort. We then tested if species’ occurrence patterns would be linked to specific traits related to the ability to cope with human presence. Finally, we assessed how road density affect the community compositional dissimilarity. We estimated 24 (41%) and 12 (21%) species to be negatively and positively affected by roads, respectively. Increased road encroachment was positively related with a higher prevalence of urban dwellers and negatively related with the occurrence of species nesting and foraging on the ground. Furthermore, increasing road density translated into an increasing community compositional dissimilarity, mostly due to species turnover. Our study support previous research showing that roads have different effects on the occurrence of different species, but we revealed that at least three species' traits have a clear relation with such road responses. Such trait filtering is probably causing a high species turnover between songbird communities occurring in roaded and nearby roadless areas. Overall, we found that different species-specific responses to roads translates into changes at the community level. Landscape and road-network management should be conceived acknowledging that roads are contributing to biodiversity changes. As so, building upon the concepts of Land Sharing / Land Sparing, conservation actions should be tailored according to the different species responses e.g., road verge management targeting species having a positive relation with road density; and compensation actions targeting species showing a negative response toward roads.

## Introduction

Roads have become the most conspicuous infrastructure in the landscapes of developed countries (Ibisch et al., 2016; Laurance and Balmford, 2013). These infrastructures are known to promote direct negative impacts on biodiversity, including roadkill, barrier and edge effects, pollution and perturbation (Trombulak and Frissell, 2000; Van der Ree et al., 2015). Such impacts may foster local population depletion and a reduction of habitat quality (Cooke et al., 2020b; D’Amico et al., 2016). Yet, roads may also benefit some species, for example by providing foraging habitat and perching structures for hunting and nesting (Ascensão et al., 2012; Morelli et al., 2014). As such, it is likely that an increase in road encroachment affect the composition of the community and, consequently, the different ecological processes and ecosystem functioning. We refer to road encroachment to describe the advancement of roads into non-urban areas, involving habitat change and increased human presence (e.g., urban sprawl and agriculture activities).

Improving our understanding on the effects associated to road encroachment on species and communities is therefore important for both road and landscape management, as well for biodiversity conservation. Yet, while we know that roads affect the occurrence of several species (Cooke et al., 2020b; D’Amico et al., 2016; Santos et al., 2016), there is still a gap of knowledge regarding how they affect the composition of ecological communities. This gap of knowledge is especially relevant given the rarity of studies on road effects focusing on large extents, from landscape to global scales, despite their value to support decisions for planning road mitigation measures (Barrientos et al., 2021). Here, we questioned whether road encroachment leads to significant changes (positive and negative) in species occurrence, and whether those changes are linked to specific traits related to perturbation sensitiveness and therefore acting as filters and increasing the community compositional dissimilarity.

We focused our study on birds, and particularly on songbirds, as these species are key components of ecosystems by providing a multitude of services, being pest predators, pollinators or seed dispersers, among others roles (Whelan et al., 2008). Also, songbirds are a widely distributed and well-studied group, especially by systematic point-count surveys, thus providing suitable data for large-scale analyses and inferences. The composition of songbird communities is also an important indicator of environmental health and ecosystem integrity (Fraixedas et al., 2020). On the other hand, human activities have greatly impacted songbird populations, with estimates of 20-25% of global reduction in the number of bird individuals from pre-agricultural times (Gaston et al., 2003), and a 27-30% decline from 1970s abundance in North America (Rosenberg et al., 2019). In Europe, there is also mounting evidence of ongoing farmland songbird decline and compositional change (Donald et al., 2001; Inger et al., 2015). Moreover, there is growing evidence that roads may be contributing to this general decline (Cooke et al., 2020b; Santos et al., 2016). Consequently, songbirds are not only suitable study species due to methodological issues, but they are also an ecologically relevant group globally threatened by roads.

We tested the following three hypotheses: *H_1_*) songbird species are differentially affected by roads, and therefore their probability of occurrence can be negatively, neutrally, or positively associated to local road density; *H_2_*) the effect of roads on species is not random, being instead linked to a prevalence of traits, overall indicating a process of filtering; and *H_3_*) such specific responses lead to changes in community composition dissimilarity. We expected to observe a higher number of species avoiding areas with higher road encroachment, given the associated threat of roadkill and increased perturbations (e.g., pollution or direct disturbance by human presence). Still, we expected species able to cope with human presence (e.g., synanthropic species) to be more prevalent in areas with a higher road density. As such, some traits related to resource exploitation and ability to cope with human activities may allow some species to take advantage of road presence, and therefore be more prevalent in areas with higher road encroachment. Furthermore, such responses were expected to increase the compositional dissimilarity when comparing areas with high and low road density, respectively. Yet, it was not easily foreseen if such increased community dissimilarity would be dominated by species turnover (i.e., species replacement) or by the loss or gain of species between roadside and roadless communities (i.e., specie nestedness).

We focused on a large and coherent biogeographically unit, the Iberian Peninsula, covering nearly 600,000 km^2^ and including regions of considerable physiographic and climatic heterogeneity. It is a discrete, relatively isolated biogeographical unit, as its connection to the European mainland is crossed by the Pyrenean Mountain chain. These characteristics make this region particularly suitable for large-scale ecological studies. Yet, our framework can easily be replicated in any other region and taxa, producing key information for planning mitigation measures.

## Methods

We analysed the data from the eBird (Sullivan et al., 2014; Wood et al., 2011), one of the most comprehensive and successful citizen-science platforms, which permitted to compile a geographically wide knowledge over a relevant ecological group. We modelled the species occurrence (presence/absence), bird traits and community compositional dissimilarity in relation to road density, while accounting for other predictors that may influence bird occurrence, namely land use and climate. We used QGIS 3.12 (QGIS.org, 2020) and R 3.6.3 (R Development Core Team, 2020) for all data preparation and analyses.

### Bird data

The citizen-science platform eBird is a web-enabled community of bird watchers that aims users to collect, manage, and store their observations in a globally accessible unified database (available in URL: ebird.org), therefore providing a large and comprehensive open dataset on bird species occurrence (Sullivan et al., 2014; Wood et al., 2011). Importantly, contributors upload the information regarding the sampling effort, including time and number of observers, survey method (e.g., point-count or walking transect), location (coordinates) and date, together with the species information, allowing data to be used in ecological research (Johnston et al., 2020). We downloaded eBird datasets for Iberian Peninsula (Portuguese and Spanish continental territories, hereafter referred as Iberia, Fig.1), for the last ten years, between 2011 and 2021.

**Figure 1.**
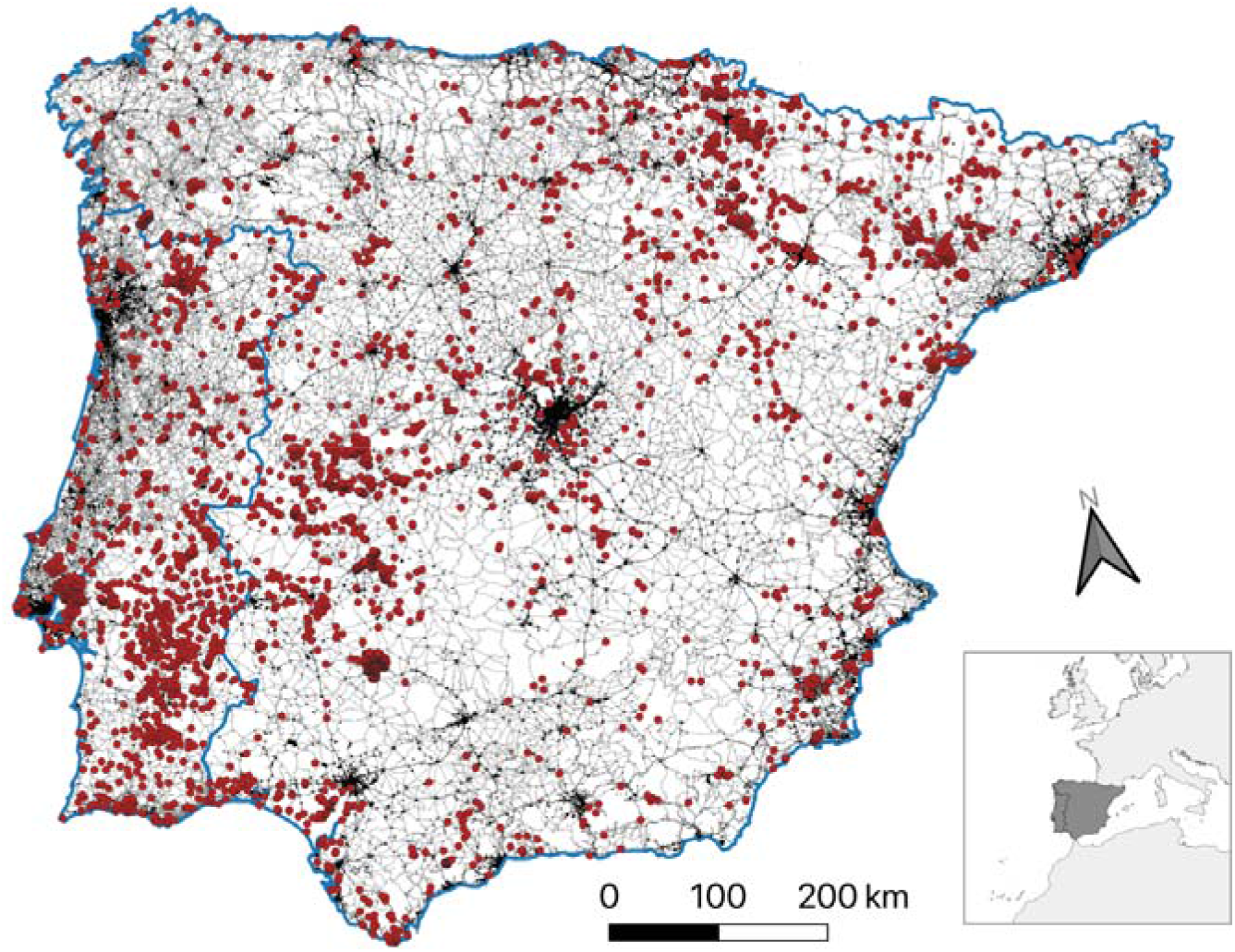
Point-count location (dots) in Iberian Peninsula (dark area in inset) used in analyses. Black areas and lines stand for major urban areas and paved roads, respectively.

### Road data

We compiled information of road spatial distribution from OpenStreetMap (OSM) using the R package ‘osmdata’ (Padgham et al., 2017). Nearly all roads have been digitized for European countries (Barrington-Leigh and Millard-Ball, 2017). We retained only the paved roads (i.e., those classified in OSM as “motorway”, “trunk”, “primary”, “secondary”, and “tertiary”), and respective ‘links’ (Fig.1), as these have medium/high traffic volume and thus are more likely to have stronger positive (i.e., attraction) or negative (e.g., population depletion or avoidance) effects on birds (Cooke et al., 2020b).

### Environmental data

We compiled information for a set of environmental variables that may influence the distribution pattern of bird species, namely land cover and altitude (TABLE 1). Environmental variables were chosen based on bibliography reporting the importance of land cover and climate as determinants of bird occurrence (Howard et al., 2020; Morelli et al., 2013). Altitude is known to correlate important climatic patterns, notably temperature and precipitation, while also affecting species distribution (Irl et al., 2015). Land cover was obtained from Copernicus’ CORINE land cover, from which we considered the nine dominant classes covering >70% of Iberian Peninsula (TABLE 1). We also aggregated all CORINE classes related to urban areas (classes 111 to 142 of CORINE), which were combined into a single Urban layer. This Urban layer was used to remove point-counts performed in urban areas (see ‘Point-count data pre-processing’ below). For each land cover class (including Urban), we created an individual layer of presence/absence. As the CORINE has a 100 m resolution and bird species are likely to be influenced by land cover at larger extents, we aggregated each individual CORINE class layer to 1×1 km resolution, in which each raster cell represents the proportion of cover (0%-100%). We used the Digital Elevation Model from Copernicus (DEM; TABLE 1) to obtain the altitude layer, which was also aggregated to 1×1km resolution (using mean values). We then ran a Principal Component Analysis on centred and scaled environmental variables (CORINE except Urban, and DEM) to reduce the multi-dimensionality and the correlations among environmental variables. We preserved the first five principal components (raster layers), which altogether retained 60% of variation, and were used in following analyses.

**TABLE 1.** Road and Environmental information considered to examine how roads affect the bird occurrence.

| Name and source | Observations |
| --- | --- |
| Road information |  |
| Paved road density; OpenStreetMap via R environment (Padgham et al., 2017) | Density of paved roads. We used a custom R code with the package ‘osmdata.’ |
| Environmental information |  |
| CORINE land cover<br>https://land.copernicus.eu/pan-european/corine-land-cover/clc2018 | Inventory of land cover with 44 classes. Year 2018. Resolution of 100 m. Main 11 classes here considered (% of cover): non-irrigated arable land (17.8), broad-leaved forest (9.9), coniferous forest (8.5), sclerophyllous vegetation (8.4), transitional woodland-shrub (6.8), natural grasslands (5.9), agro-forestry areas (5.4), permanently irrigated land (5.1), olive groves (4.6), overall covering >70% of Iberia. All urban related classes were aggregated into a dingle Urban class. |
| Digital Elevation Model<br>https://land.copernicus.eu/imagery-in-situ/eu-dem/eu-dem-v1.1 | Year 2016. Resolution of 25 m. |

### Data analysis

#### Point-count data pre-processing

We filtered the eBird data keeping only the complete checklists (a complete checklist is any eBird list where birding is the primary purpose, and every species is identified to the best of surveyor ability), performed using the protocol “stationary count” (point-counts), with the duration of 5 to 15 minutes, and by up to four observers. Most point-counts in our dataset (87% of total) were performed by a single observer for 5 min (43%), 5-10 min (27%) and 10-15 min (17%).

We restricted our analysis to most songbirds, as these species are more likely to be recorded and targeted for while performing point-count surveys. As such, we retained all *Passeriformes* except those species from the families *Laniidae*, *Corvidae*, *Cinclidae*, and *Hirundinidae* (species from these four families are not usually monitored in point-counts). As the bird community changes significantly between breeding and non-breeding seasons, and roads probably affect resident species differently during these periods (e.g., when choosing nesting sites), we opted to filter the data to consider only point-counts performed during the breeding season (from March to June). We further filtered the dataset to include surveys performed during the morning period (between 5:00 and 11:59 am), as in this period all songbirds are more active and therefore more likely to be detected. This pre-processing allowed us to keep surveys with known survey effort and performed with a relatively similar sampling protocol across the entire study area. It also provided high confidence that the records reported were obtained on the location reported (point-count), reducing the probability of being recorded during a travelling census (i.e., crossing different habitats and distance to putative roads).

Each retained point-count site was then characterized according to the five principal components of environmental information by extracting the corresponding value of each layer at the survey site. We discarded those point-counts performed in areas with > 20% of Urban, as bird communities inhabiting more dense urban areas are exposed to several human-related pressures (Callaghan et al., 2019; Maklakov et al., 2011; McKinney, 2006) which, together with the existing higher density of roads, could potentially confound the effects of road encroachment. Also, as the detectability of birds can be affected by the proximity of roads with high traffic volumes (Cooke et al., 2020a), we further discarded those point-counts carried out close (<50 m) to a main road (highway, trunk and primary, and respective links). There is a low effect of low frequency background noise (traffic) on the surveyors’ ability to detect birds at >50 m (Pacifici et al., 2008).

For each remaining point-count site, we calculated the road density by summing all paved road segments intercepting a 1 km buffer radius from the point-count site. We choose this distance assuming that most significant road-effect zones span within 1 km from paved road (Forman and Deblinger, 2000). For each point-count event, we calculated the survey effort as number of observers x survey time. Our dataset comprised 112 songbird species, recorded during 9,397 survey events in 4,459 unique point-count sites (see TABLE S1.1 in Supplementary material S1 for species names and totals).

#### Road effects on species occurrence

To test if species are differentially affected by roads (*H_1_*), we assessed the relationship between the species’ presence/absence and the road density and environmental predictors, using Generalized Linear Models (GLM) with binomial error and logit link function. Road density was scaled to allow comparisons between models estimates for this variable. As one third of the point-count sites were surveyed on more than one occasion, we aggregated the information from the different surveys retaining all registered species (presences) and summing all survey effort. Total survey effort was included in GLMs as an offset (log-transformed). Importantly, for each species, we only considered those point-counts performed where the species is resident year-round or breeds, which was assured by overlaying the point-count locations (with presences and absences) with the corresponding information from the species distribution as compiled by the BirdLife International (data retrieved in May 2021). Before analyses, we searched for collinearity between road, environmental information (the five principal components), and survey effort using Pearson correlations. All pairs of variables had Pearson’s |*r*| < 0.6 and therefore no variable was discarded.

We only considered those species recorded in at least 50 distinct point-count sites to guarantee sufficient model discrimination power (see TABLE S1.1). Model discrimination power was assessed by the area under the receiver operational curve (AUC). For each species, the coefficient of the Road variable and its 95% confidence interval (CI) of the respective GLM were used to classify the species with respect to its response to roads as follows: as ‘negatively’ or ‘positively’ associated with road density if the CI was below/above zero, respectively, and as ‘neutral’ if zero was within the CI.

#### Road effects on functional diversity

To test if the species’ occurrence patterns would be linked to specific traits related to the ability to cope with human presence and activities (*H_2_*), we considered six ecological traits corresponding to major differences in bird life strategy, grouped into three main trait categories: *Physical attributes*, *Environmental tolerance,* and *Resource use*.

For *Physical attributes*, we analysed the body mass as this trait is related to many others physical attributes, namely height or wingspan. We expected that larger species would be less frequent in areas with higher road encroachment, as many of them are more sensitive to roadkill and consequent population depletion (Santos et al., 2016). We also analysed the brain mass, using the residuals of the linear model relating brain volume and body mass, given the high correlation between both traits (Pearson’s *r*>0.8). It has been shown that brain mass correlate with several measures of behavioural flexibility and ability to survive in novel ecological conditions (Callaghan et al., 2019; Fristoe et al., 2017; Maklakov et al., 2011). We therefore expected a positive association between road encroachment and brain mass.

For traits related to *Environmental tolerance*, we analysed the ability to thrive in cities (city dwelling or urban tolerance). We expected road encroachment to be related to an increase of city dwellers in rural areas (McKinney, 2006). We further tested if roads could filter species nesting on the ground, expecting that higher road encroachment refrain such species to occur in the breeding season, given their vulnerability to nest predation, particularly of domestic cats associated to humans (Pita et al., 2009).

Finally, for *Resource use*, we analysed the diet and the foraging behaviour. Birds feeding on invertebrates are likely to have higher visual acuity and therefore be able to better perceive incoming vehicles. Also, road verges may have vegetated areas with low management intensity, particularly when compared to intensive agricultural areas (Ascensão et al., 2012), thus benefiting insectivorous birds due to higher insect abundance (Muñoz et al., 2015; Villemey et al., 2018). Hence, we expect a higher prevalence of invertivorous species with increasing road encroachment. Likewise, the foraging behaviour was expected to affect species’ responses to road encroachment, with those species spending more time foraging at the ground level avoiding higher road encroachment due to higher human activity therein.

For each individual trait considered, we built different phylogenetic regression models, relating the coefficient of the road variable obtained in the GLM as the response variable and the trait value per species as single predictor. Phylogenetic regression models allow controlling for phylogenetic relatedness, as related species tend to share many traits due to shared evolutionary history, and therefore cannot be used in statistical analysis as truly independent observations (Paradis et al., 2004). For body mass, brain mass (log transformation), diet (arcsin transformation), and foraging (arcsin transformation), we used phylogenetic linear models; for ground nesting and urban tolerance we used phylogenetic logistic regression models. We only considered those species previously classified as being ‘negatively’ or ‘positively’ associated to roads to reduce the nuisance from ‘neutral’ responsive species, and therefore increase the chance of detecting significant differences.

We replicated the phylogenetic regression models, for each trait, across 100 random and equally plausible phylogenetic trees (replicates) to evaluate the consistency of the relation between road response and each trait, as a sensitivity analysis. The distributions of the 95% confidence intervals of the regression coefficients of each trait, per replicate, were used to classify the association of the road responsive value and each trait (positive, negative, or neutral), per replicate. Phylogenetic trees were based on the Ericson backbone (Ericson et al., 2006) and obtained from the BirdTree (Jetz et al., 2012). Phylogenetic regressions were performed using the R package ‘phylolm’ (Tung Ho and Ané, 2014) (see Supplementary material S2 on phylogenetic regression models and compiled trait data).

#### Road effects on community dissimilarity

To test the effect of roads on community dissimilarity (*H_3_*), we used the full dataset (112 species). We selected paired point-counts, in which each pair had one point-count performed in a roadless area (i.e., no paved roads in a radius of 1 km), and one point-count nearby having paved roads within that buffer radius. The selection of pairs of points obeyed to the following sequential rules: first, for each roadless point-count (randomly selected), we searched for a roaded point-count nearby i.e., within 1100 m radius. If more than one neighbour existed, we randomly selected one site. If there was no neighbouring site, we increased the search radius by 100 m. This procedure was repeated until at least one neighbouring roaded point-count was selected or the search radius reached a maximum of 2000 m, so that the roadless point-count location would be beyond the road-effect zone distance while ensuring similar environmental conditions. Each point-count location was used in only one pair of points, to reduce pseudo-replication issues.

There are two potential ways in which road and roadless species assemblages can be ‘different’: by species replacement (i.e., turnover), with the substitution of species in one site by different species in the other site; and by species loss or gain, which implies the elimination or addition of species in only one of the sites, and leads to the poorest assemblage being a strict subset of the richest one (i.e., nestedness) (see Baselga, 2010). We applied this approach for partitioning the total Sørensen dissimilarity into the two components, obtaining the dissimilarity derived solely from turnover and the dissimilarity derived from nestedness, allowing us to assess which process dominates eventual changes in communities. Dissimilarity metrics were calculated using the R package ‘betapart’ (Baselga and Orme, 2012)

We further used the total Sørensen dissimilarity and their components (turnover and nestedness) as response variables in three beta regression models. The difference in road density across each pair of point-counts was used as a predictor. To further control for environmental and survey effort differences across the pairwise points, the models also integrated information on *i*) the Euclidean distance between sites; *ii*) the distance between the five PCA components, and *iii*) the difference in survey effort. The distance between the five PCA components were computed as 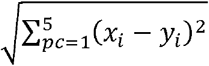, where *pc* are the principal components. Beta regression models were built using the R package ‘glmmTMB’ (Brooks et al., 2017)

## Results

### Road effects on species occurrence

We built occurrence models for 58 focal species, which achieved a mean±SD AUC of 0.73±0.05 (range: 0.64-0.86), representing a reasonable discrimination power across the models. More than one-third of these species (n=24, 41%) were classified as being negatively affected by roads (such as the Dartford warbler *Sylvia undata*, globally listed as Near Threatened by the IUCN), while 12 (21%) were classified as being positively affected by roads (such as the Eurasian blackbird *Turdus merula*), and 22 (38%) as neutral (Fig.2).

**Figure 2.**
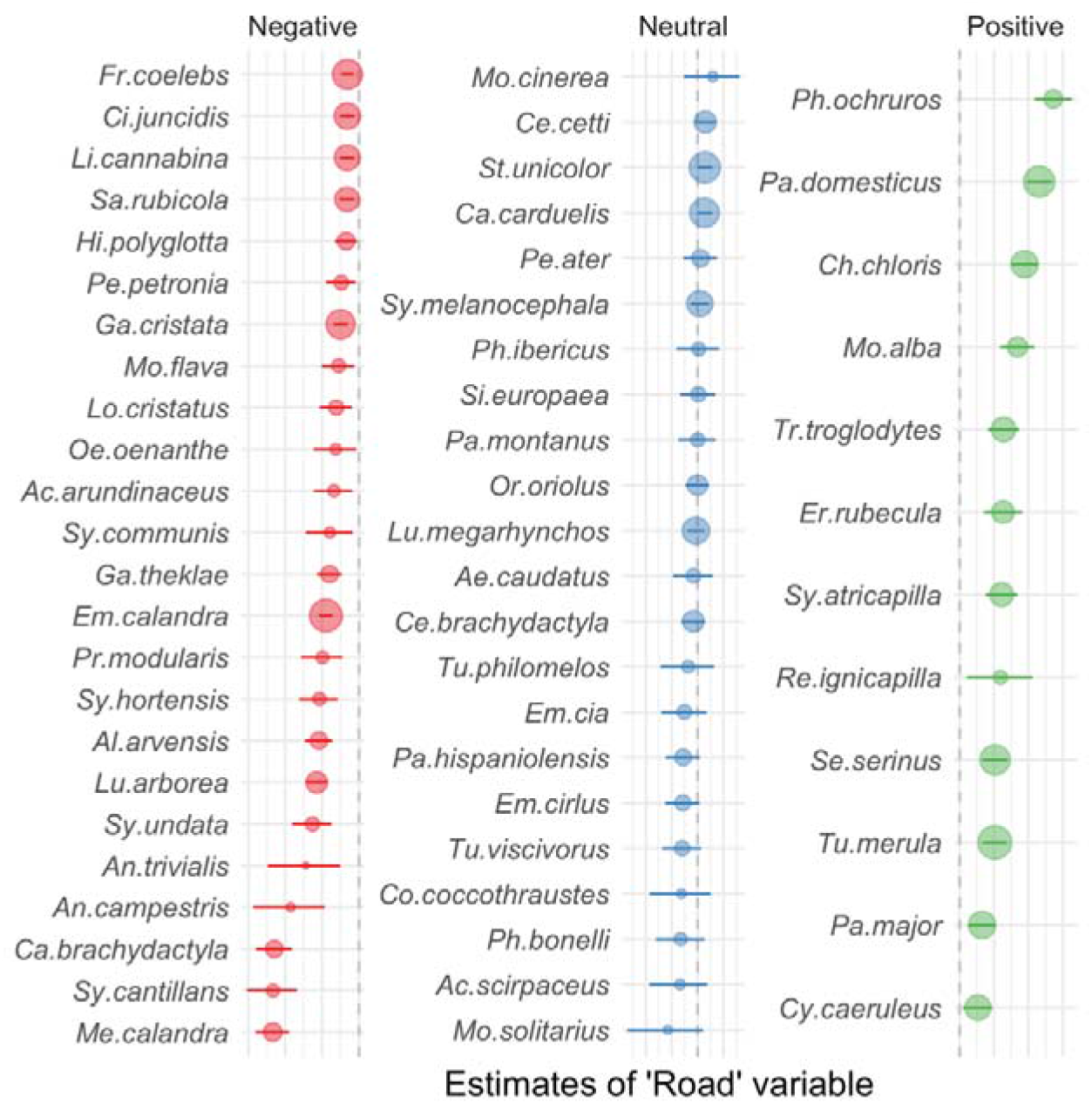
Association type (Negative, Neutral, or Positive) between road density and species occurrence. The association was determined by the estimate and confidence intervals of the Road variable from the GLMs relating species occurrence in point-counts with road and environmental variables. Size of the dot is proportional to the number of point-count sites in which the species was recorded. Vertical dashed line represents zero value. Complete scientific names of species are shown in Table S1.1.

### Road effects on functional diversity

The individual phylogenetic models suggest that increased road encroachment is positively related with a higher prevalence of urban dwellers and negatively related with the occurrence of species nesting on ground and foraging on the ground (the proportion of replicates having CI overlapping zero were 0%, 4% and 11%, respectively; Fig.3C, D, F). Results further suggest that road encroachment may be somehow positively associated with higher brain mass, as all the estimates across replicates were positive and away from zero (although 81% of CI of replicates crosse zero; Fig. 3B). There was no evidence for an association of body mass of invertivory with road density (Fig. 3A, E).

**Figure 3.**
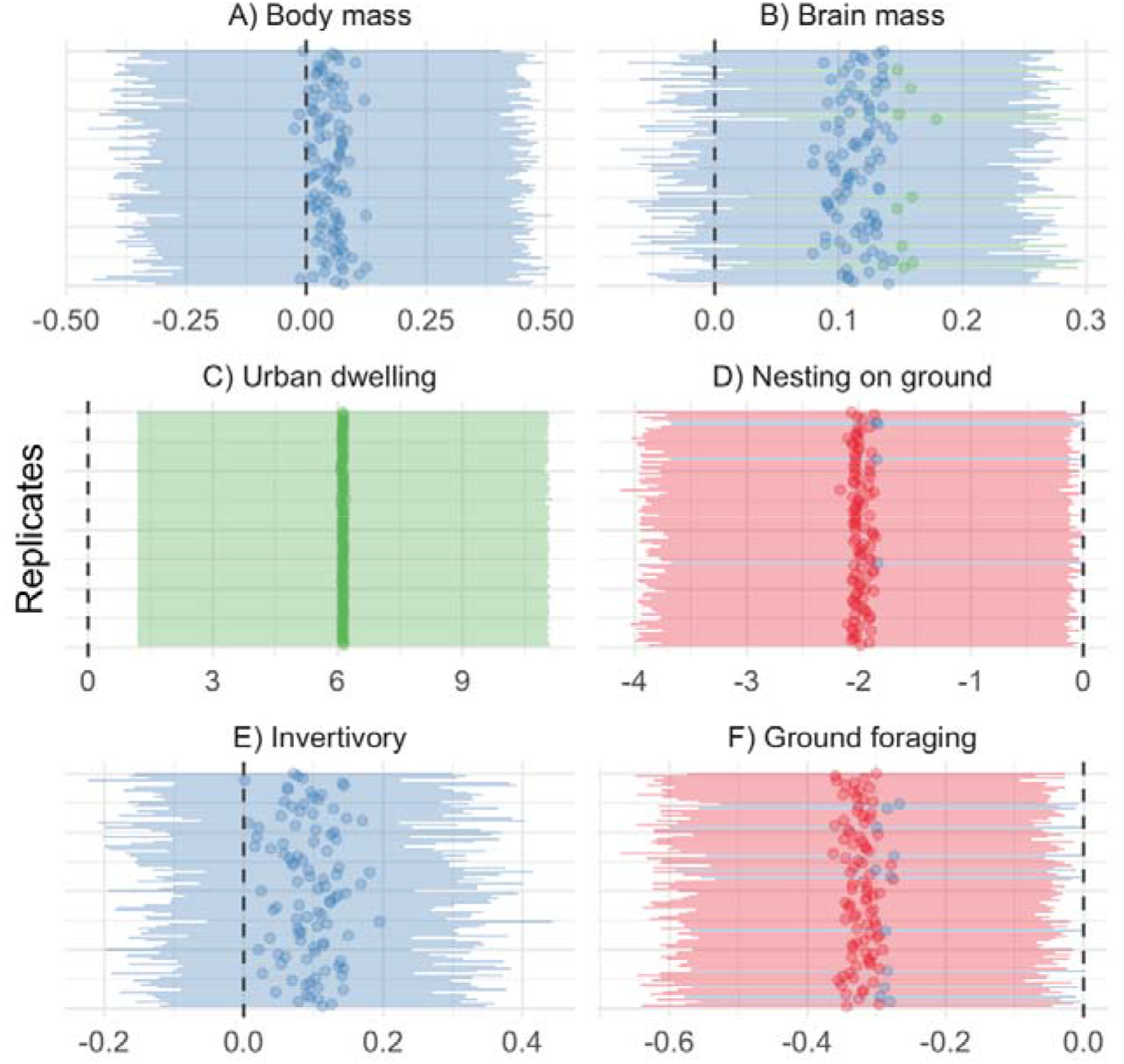
Relation between road density and six traits associated to physical attributes (A, B), environmental tolerance (C, D), and resource use (E, F). Each line stands for one replicate of phylogenetic regression models using different equally plausible phylogenetic trees. The dots and bars indicate the estimate and CI of the mean road density variable, coloured according to the type of relation with the trait (negative, not significant, and positive).

As an outcome of the phylogenetic models, we searched for ‘super-tolerant’ and ‘super-avoider’ species towards roads. Super-tolerant species were those bringing together all the traits related to road use i.e., city dwellers not nesting on the ground and with null or low foraging on the ground. Two species previously classified as being positively associated with roads showed all these traits, namely the great tit *Parus major* and blue tit *Cyanistes caeruleus*, but several other species (among them several warblers) showed at least two traits. On the other hand, super-avoiders were all the species bringing together all the traits related to road avoidance i.e., species inhabiting natural areas and both nesting and foraging on the ground, such as the Eurasian skylark *Alauda arvensis*, but also other lark species (Fig. 4).

**Figure 4.**
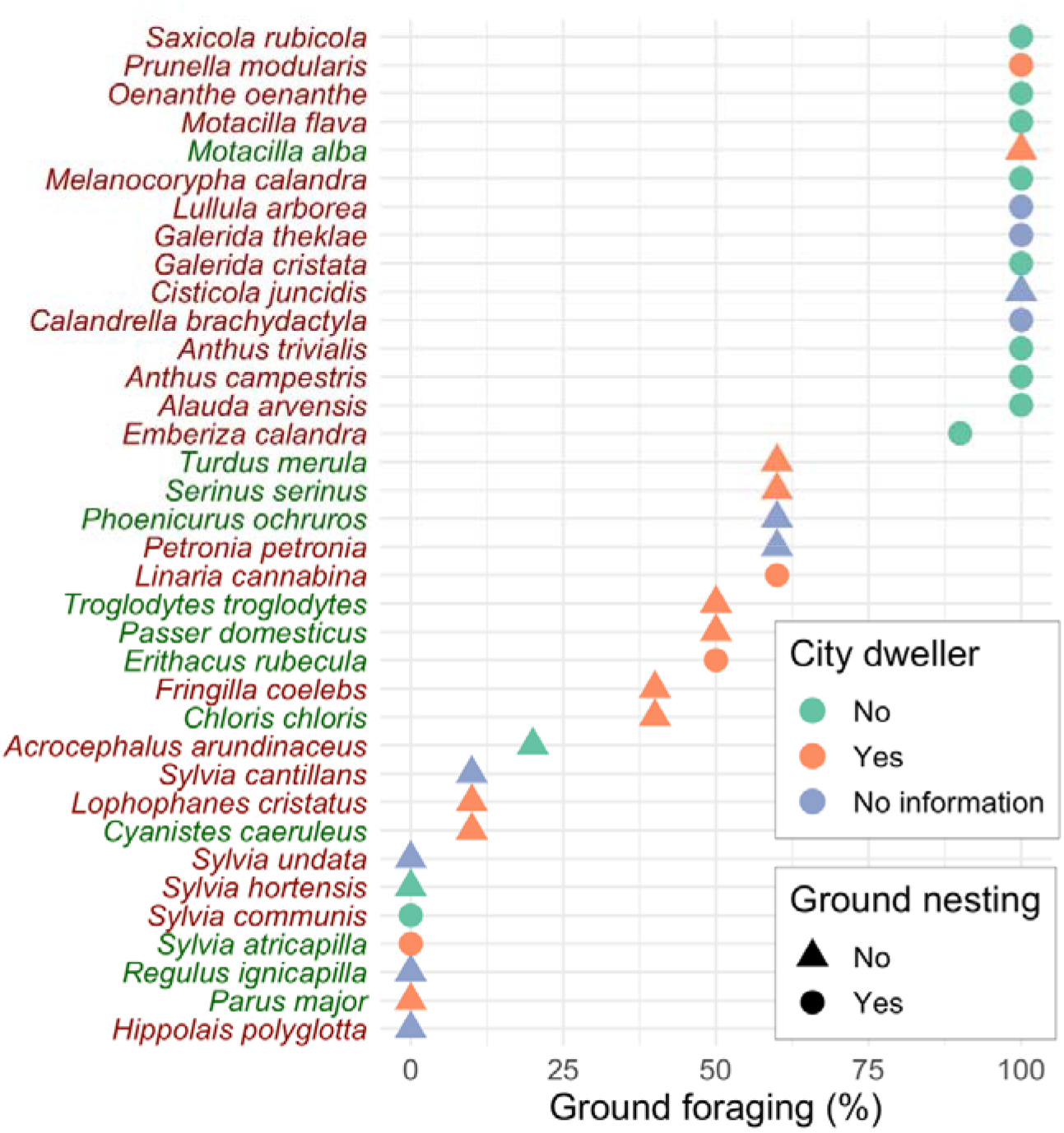
Characterization of species according to three traits that varied significantly with road density: city dwelling (symbol colours), ground nesting (symbol shape) and ground foraging (symbol position along the XX-axis). Each species is coloured according to the association between road density and species occurrence, as depicted in Fig. 2: red/green labels refer to species having a negative/positive association with road density, respectively. See Supplementary material S2 for species information.

### Road effects on community dissimilarity

We combined 311 pairs of point-counts to access the road effects on the pairwise dissimilarity (pairwise Euclidean distance averaged 1230±496 m). The mean±SD of the total Sørensen dissimilarity was 0.55±0.22. When decomposing the total dissimilarity into the turnover and nestedness components, results suggest that most of observed dissimilarity is due to turnover of species between roaded and nearby roadless sites (Fig. 5).

**Figure 5.**
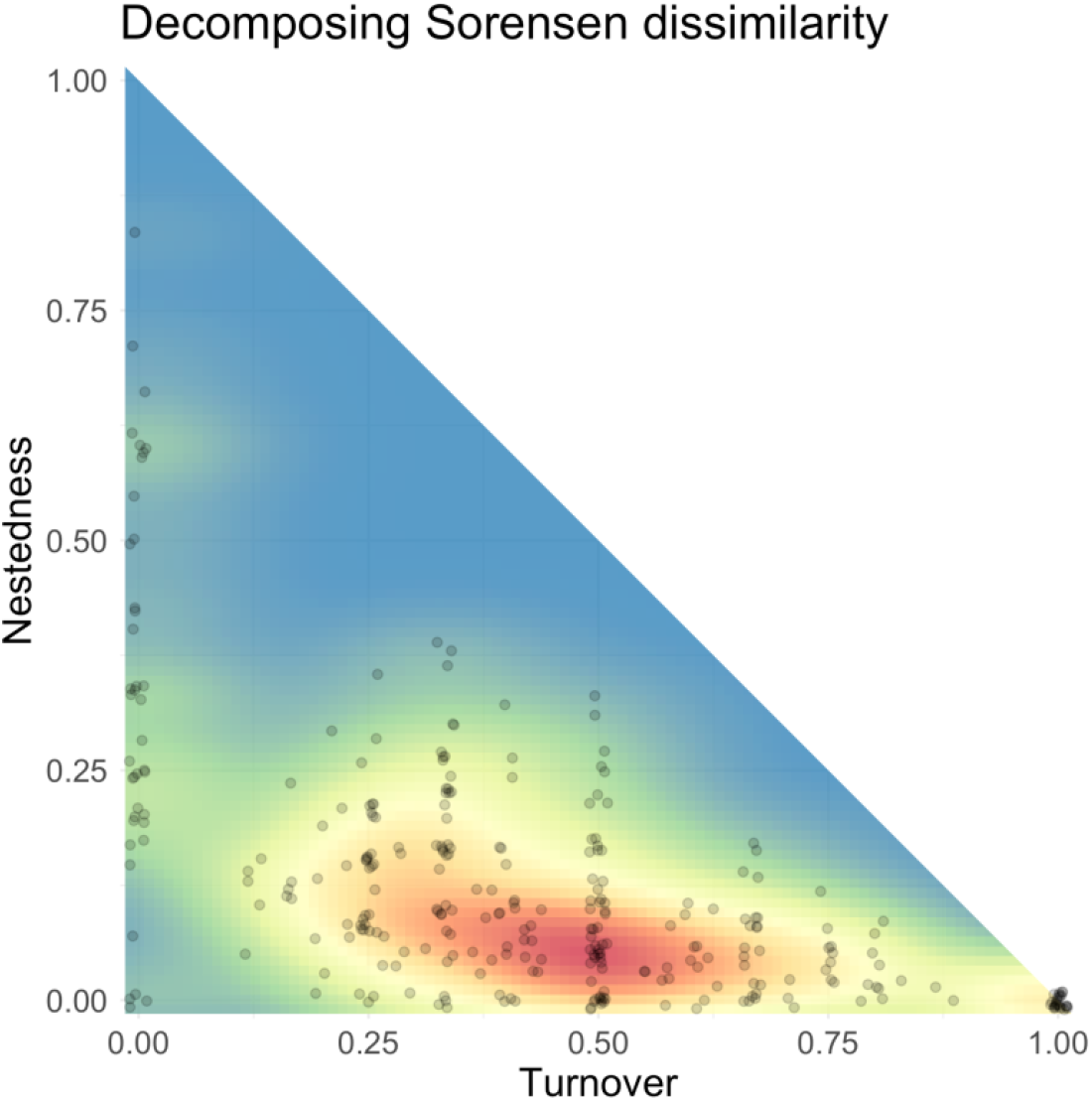
Sorensen dissimilarity decomposed into the two components turnover and nestedness. Each point stands for one pair of point count sites used to access the community dissimilarity. Colour gradient is the two-dimensional kernel density estimation, with redder colours indicating higher concentration of points.

As expected, the Sørensen dissimilarity increases with survey effort, distance between point-count sites, and with the difference in road density (TABLE 2). The turnover of species follows the same pattern, but this fold the distance between point-count sites and the difference in road density seems to be more important (TABLE 2). The component nestedness of Sørensen dissimilarity was not significantly related to any predictor (TABLE 2).

**TABLE 2.** Summary results for the beta regression models relating the community dissimilarity (total Sørensen dissimilarity and its components turnover and nestedness), with pairwise difference of environment (Env. dist.), survey effort, Euclidean distance, and road density. Est. – estimate, CI – confidence interval. * *p*<0.05 ** *p*<0.01 *** *p*<0.001.

| *Predictors* | **Total Sørensen** |  | **Turnover** |  | **Nestedness** |  |
| --- | --- | --- | --- | --- | --- | --- |
|  | *Est.* | *CI* | *Est.* | *CI* | *Est.* | *CI* |
| Intercept | **1.23** *** | **1.09 – 1.39** | **0.57** *** | **0.49 – 0.66** | **0.12** *** | **0.10 – 0.15** |
| Env. dist. | 1.08 | 0.96 – 1.22 | 0.89 | 0.77 – 1.03 | 1.11 | 0.99 – 1.24 |
| Survey effort | **1.16** * | **1.03 – 1.30** | 1.06 | 0.93 – 1.20 | 0.95 | 0.85 – 1.06 |
| Euclidean dist. | **1.13** * | **1.00 – 1.28** | **1.36** *** | **1.18 – 1.57** | 0.89 | 0.79 – 1.01 |
| Road density | **1.17** * | **1.04 – 1.32** | **1.42** *** | **1.23 – 1.64** | 0.97 | 0.86 – 1.08 |

## Discussion

We studied the responses of songbirds to the presence of roads by formulating hypotheses about the effect of road density on their occurrence, trait prevalence, and community dissimilarity, allowing to understand how changes at the species level translate into changes at community level. Our results provided evidence that roads affect species in different ways, but such effect is not random, consequently shaping the community composition over broad spatial scales. Our study support previous research showing that roads have negative, neutral and positive effects on the occurrence of different species (Cooke et al., 2020b), but we showed that at least three species' traits have a clear relation with such road responses, and that such trait filtering is probably causing a high species turnover between songbird communities occurring in roaded and nearby roadless areas.

We showed that species more sensitive to perturbation, like those nesting and foraging on the ground such as the Eurasian skylark, avoid areas with higher road density, probably due to the higher human presence and activity (Summers et al., 2011), which reduce the habitat quality for those birds and consequently their occurrence. With a massive long-term decline of ground-nesting songbird species across Europe (McMahon et al., 2020), it is possible that road encroachment and related impacts are contributing to this trend by further reducing or degrading existing habitat. Conversely, the city dwellers, such as the house sparrow *Passer domesticus,* showed a strong association with road density. These species are often generalists, known to be the most urban‐tolerant and adaptable of birds (Callaghan et al., 2019; Evans et al., 2011; McKinney, 2006). This result suggests that increasing road encroachment may promote the prevalence of urban-tolerant species that are less likely to be present in roadless areas, while impacting more sensitive species. The prevalence of city dwellers had been previously shown for urban environments, including the consequent homogenization of communities (Clergeau et al., 2006; McKinney, 2006; Proppe et al., 2013), and we suggest that this effect may spread to rural and even more natural environments as road encroachment increases. The phylogenetic models also suggested that species with larger brain mass are more likely to cope with increased road encroachment. This is probably due to their behavioural flexibility and ability to survive in novel ecological conditions (Callaghan et al., 2019; Fristoe et al., 2017; Maklakov et al., 2011). Our study further showed that the different species-specific responses to roads translate into changes at the community level, mostly due to species turnover, as previously suggested by a meta-analysis on this topic (Kroeger et al., 2021).

Such species and community-level changes may come with negative consequences for the ecosystem functioning and the provision of ecosystem services, including a high toll for economies. If so, increasing road densities may be affecting the ecosystem functioning and their integrity (Glennon and Porter, 2005), as well their ecological networks, particularly on species interactions (Tylianakis et al., 2008). For example, functions performed by certain species may become over or underrepresented, and ecosystem services may be impoverished or lost, with further costs to human activities (McWilliams et al., 2019). Note that we did not analyse data collected in urban environments, where the processes of changes are likely to be more pronounced due to the higher road densities and higher human activities therein. Likewise, we focused on paved roads, although there is some evidence that dirt roads may also promote habitat degradation with consequent impacts for some species (Cooke et al., 2020b; Mammides et al., 2016).

### Management recommendations

Landscape and road-network management should be conceived acknowledging that roads are contributing to biodiversity changes. As so, building upon the concepts of Land Sharing (i.e., the integration of nature conservation into more anthropized areas) and Land Sparing (i.e., the separation between areas planned for human development and nature conservation) (Fischer et al., 2014), we suggest that management actions should be tailored according to the different species responses.

Concerning land sharing, the management of road verges and surrounding areas should target the conservation of species having a positive relation with road density, especially super-tolerant species such as the great tit and more generalist warblers. Road verges have been already acknowledged as important areas for biodiversity conservation (Meunier et al., 1999; Phillips et al., 2020). We suggest that roadsides can play a role as refuges, also for songbirds, and recommend verges to be managed favouring the creation of habitat for super-tolerant species, especially those species providing ecosystem services. This goal can be achieved especially on wide roadsides, where there is room for creating different vegetation density and structure outside the mowed vegetation strip adjacent to the roadway. For example, a possible roadside design can include some patches with taller shrubs and other patches without shrubs but with a higher grass layer. Nevertheless, the roadkill risk should be carefully considered before promoting species presence along roadsides. For example, these actions can be primarily implemented along minor roads (i.e., having lower roadkill risk) or along major highways (i.e., generally with wide verges that allow vegetation improvements far from traffic lanes). Finally, roadkill is not the only risk related to roads, so all the possible threats (such as pollution, poaching, etc.) should be considered before promoting the presence of songbirds along roadsides.

Concerning land sparing, compensation actions (namely preserving areas devoid of roads) should target species showing a negative response toward roads, especially threatened super-avoiders. In fact, roadless areas have been targeted as key areas for conservation (Ibisch et al., 2016), and therefore should be kept free from road encroachment. However, roadless-areas' preservation should not only focus on larger protected areas. We showed that several small roadless areas are (heterogeneously) distributed across two developed countries (i.e., Spain and Portugal), and that they host relatively well-conserved songbirds’ communities. Any landscape planning should therefore consider the preservation of these biodiversity refuges. One of possible strategies to reach this goal is the optimization of road-networks, which should aim to reduce road redundancy by avoiding the existence of several alternative paths between the same urban areas, combining human connectivity into fewer transport corridors, more effective and probably multimodal. While this approach may have huge benefits for conservation and society in developing countries (Hopcraft et al., 2015), in more developed regions such optimization can be achieved through the decommissioning of redundant roads (D’Amico et al., 2016).

In all cases, the connectivity between habitat patches should be guaranteed, namely by improving the gap permeability, both along roads for tolerant species and across roads for avoiders. This can be achieved by improving the habitat quality for tolerant species along major roads and field margins, but also implementing vegetated wildlife road-crossing structures (the so-called ecoducts or overpasses) for connecting habitat patches for both tolerant species and, importantly, super-avoiders. Finally, we tested our hypotheses on a very large scale (i.e., the Iberian Peninsula), suggesting that such patterns (and related management implications) can be general in regions with similar road-network development, as in many developed countries. However, our results can also be considered preventively for road-network planning in developing countries, where a large number of new infrastructures are being planned and even built (Ascensão et al., 2018) with potential alarming implications for biodiversity conservation.

## Authors’ contributions

FA collected the data, developed the methods, performed the data analysis, and prepared the first draft of the paper. FA, MD, ER and HMP contributed to the design of the research, discussed data, and contributed to writing the final manuscript.

## Acknowledgements

We thank all eBird contributors, without whom research such as this would not be possible, and the eBird program for providing this platform and facilitating data analyses. FA was funded by Fundação para a Ciência e Tecnologia (contract CEECIND/03265/2017). MD was funded by a JdC-Inc postdoctoral grant (IJC2019-039662-I) from the Spanish Ministry of Science and Innovation (MICINN). The authors declare that there is no conflict of interest.

## References

Ascensão, F., Clevenger, A.P., Grilo, C., Filipe, J., Santos-Reis, M., 2012. Highway verges as habitat providers for small mammals in agrosilvopastoral environments. Biodivers. Conserv. 21, 3681–3697. https://doi.org/10.1007/s10531-012-0390-3

Ascensão, F., Fahrig, L., Clevenger, A.P., Corlett, R.T., Jaeger, J.A.G., Laurance, W.F., Pereira, H.M., 2018. Environmental challenges for the Belt and Road Initiative. Nat. Sustain. 1, 206–209. https://doi.org/10.1038/s41893-018-0059-3

Barrientos, R., Ascensão, F., D’Amico, M., Grilo, C., Pereira, H.M., 2021. The lost road: do transportation networks imperil wildlife population persistence? Perspect. Ecol. Conserv. https://doi.org/10.1016/j.pecon.2021.07.004

Barrington-Leigh, C., Millard-Ball, A., 2017. The world’s user-generated road map is more than 80% complete. PLOS ONE 12, e0180698. https://doi.org/10.1371/journal.pone.0180698

Baselga, A., 2010. Partitioning the turnover and nestedness components of beta diversity: Partitioning beta diversity. Glob. Ecol. Biogeogr. 19, 134–143. https://doi.org/10.1111/j.1466-8238.2009.00490.x

Baselga, A., Orme, C.D.L., 2012. betapart: an R package for the study of beta diversity. Methods Ecol. Evol. 3, 808–812. https://doi.org/10.1111/j.2041-210X.2012.00224.x

Brooks, M., E., Kristensen, K., Benthem, K., J., van, Magnusson, A., Berg, C., W., Nielsen, A., Skaug, H., J., Mächler, M., Bolker, B., M., 2017. glmmTMB Balances Speed and Flexibility Among Packages for Zero-inflated Generalized Linear Mixed Modeling. R J. 9, 378. https://doi.org/10.32614/RJ-2017-066

Callaghan, C.T., Major, R.E., Wilshire, J.H., Martin, J.M., Kingsford, R.T., Cornwell, W.K., 2019. Generalists are the most urban-tolerant of birds: a phylogenetically controlled analysis of ecological and life history traits using a novel continuous measure of bird responses to urbanization. Oikos 0. https://doi.org/10.1111/oik.06158

Clergeau, P., Croci, S., Jokimäki, J., Kaisanlahti-Jokimäki, M.-L., Dinetti, M., 2006. Avifauna homogenisation by urbanisation: Analysis at different European latitudes. Biol. Conserv. 127, 336–344. https://doi.org/10.1016/j.biocon.2005.06.035

Cooke, S.C., Balmford, A., Johnston, A., Massimino, D., Newson, S.E., Donald, P.F., 2020a. Road exposure and the detectability of birds in field surveys. Ibis 162, 885–901. https://doi.org/10.1111/ibi.12787

Cooke, S.C., Balmford, A., Johnston, A., Newson, S.E., Donald, P.F., 2020b. Variation in abundances of common bird species associated with roads. J. Appl. Ecol. 57, 1271–1282. https://doi.org/10.1111/1365-2664.13614

D’Amico, M., Périquet, S., Román, J., Revilla, E., 2016. Road avoidance responses determine the impact of heterogeneous road networks at a regional scale. J. Appl. Ecol. 53, 181–190. https://doi.org/10.1111/1365-2664.12572

Donald, P.F., Green, R.E., Heath, M.F., 2001. Agricultural intensification and the collapse of Europe’s farmland bird populations. Proc. R. Soc. Lond. B Biol. Sci. 268, 25–29. https://doi.org/10.1098/rspb.2000.1325

Ericson, P.G., Anderson, C.L., Britton, T., Elzanowski, A., Johansson, U.S., Källersjö, M., Ohlson, J.I., Parsons, T.J., Zuccon, D., Mayr, G., 2006. Diversification of Neoaves: integration of molecular sequence data and fossils. Biol. Lett. 2, 543–547.

Evans, K.L., Chamberlain, D.E., Hatchwell, B.J., Gregory, R.D., Gaston, K.J., 2011. What makes an urban bird? Glob. Change Biol. 17, 32–44. https://doi.org/10.1111/j.1365-2486.2010.02247.x

Fischer, J., Abson, D.J., Butsic, V., Chappell, M.J., Ekroos, J., Hanspach, J., Kuemmerle, T., Smith, H.G., Wehrden, H. von, 2014. Land Sparing Versus Land Sharing: Moving Forward. Conserv. Lett. 7, 149–157. https://doi.org/10.1111/conl.12084

Forman, R.T.T., Deblinger, R.D., 2000. The Ecological Road-Effect Zone of a Massachusetts (U.S.A.) Suburban Highway. Conserv. Biol. 14, 36–46. https://doi.org/10.1046/j.1523-1739.2000.99088.x

Fraixedas, S., Lindén, A., Piha, M., Cabeza, M., Gregory, R., Lehikoinen, A., 2020. A state-of-the-art review on birds as indicators of biodiversity: Advances, challenges, and future directions. Ecol. Indic. 118, 106728. https://doi.org/10.1016/j.ecolind.2020.106728

Fristoe, T.S., Iwaniuk, A.N., Botero, C.A., 2017. Big brains stabilize populations and facilitate colonization of variable habitats in birds. Nat. Ecol. Evol. 1, 1706. https://doi.org/10.1038/s41559-017-0316-2

Gaston, K.J., Blackburn, T.M., Goldewijk, K.K., 2003. Habitat conversion and global avian biodiversity loss. Proc. R. Soc. Lond. B Biol. Sci.

Glennon, M.J., Porter, W.F., 2005. Effects of land use management on biotic integrity: An investigation of bird communities. Biol. Conserv. 126, 499–511. https://doi.org/10.1016/j.biocon.2005.06.029

González-Suárez, M., Zanchetta Ferreira, F., Grilo, C., 2018. Spatial and species-level predictions of road mortality risk using trait data. Glob. Ecol. Biogeogr. 27, 1093–1105. https://doi.org/10.1111/geb.12769

Hopcraft, J.G.C., Mduma, S.A.R., Borner, M., Bigurube, G., Kijazi, A., Haydon, D.T., Wakilema, W., Rentsch, D., Sinclair, A.R.E., Dobson, A., Lembeli, J.D., 2015. Conservation and economic benefits of a road around the Serengeti. Conserv. Biol. 29, 932–936. https://doi.org/10.1111/cobi.12470

Howard, C., Stephens, P.A., Pearce□Higgins, J.W., Gregory, R.D., Butchart, S.H.M., Willis, S.G., 2020. Disentangling the relative roles of climate and land cover change in driving the long-term population trends of European migratory birds. Divers. Distrib. 26, 1442–1455. https://doi.org/10.1111/ddi.13144

Ibisch, P.L., Hoffmann, M.T., Kreft, S., Pe’er, G., Kati, V., Biber-Freudenberger, L., DellaSala, D.A., Vale, M.M., Hobson, P.R., Selva, N., 2016. A global map of roadless areas and their conservation status. Science 354, 1423–1427. https://doi.org/10.1126/science.aaf7166

Inger, R., Gregory, R., Duffy, J.P., Stott, I., Voříšek, P., Gaston, K.J., 2015. Common European birds are declining rapidly while less abundant species’ numbers are rising. Ecol. Lett. 18, 28–36. https://doi.org/10.1111/ele.12387

Irl, S.D.H., Harter, D.E.V., Steinbauer, M.J., Puyol, D.G., Fernández-Palacios, J.M., Jentsch, A., Beierkuhnlein, C., 2015. Climate vs. topography – spatial patterns of plant species diversity and endemism on a high-elevation island. J. Ecol. 103, 1621–1633. https://doi.org/10.1111/1365-2745.12463

Jetz, W., Thomas, G., Joy, J., Hartmann, K., Mooers, A., 2012. The global diversity of birds in space and time. Nature 491, 444.

Johnston, A., Hochachka, W.M., Strimas-Mackey, M.E., Gutierrez, V.R., Robinson, O.J., Miller, E.T., Auer, T., Kelling, S.T., Fink, D., 2020. Analytical guidelines to increase the value of citizen science data: using eBird data to estimate species occurrence. bioRxiv 574392. https://doi.org/10.1101/574392

Kroeger, S.B., Hanslin, H.M., Lennartsson, T., D’Amico, M., Kollmann, J., Fischer, C., Albertsen, E., Speed, J.D.M., 2021. Impacts of roads on bird species richness: A meta-analysis considering road types, habitats and feeding guilds. Sci. Total Environ. 151478. https://doi.org/10.1016/j.scitotenv.2021.151478

Laurance, W.F., Balmford, A., 2013. A global map for road building. Nature 495, 308–309. https://doi.org/10.1038/495308a

Maklakov, A.A., Immler, S., Gonzalez-Voyer, A., Rönn, J., Kolm, N., 2011. Brains and the city: big-brained passerine birds succeed in urban environments. Biol. Lett. 7, 730–732.

Mammides, C., Kounnamas, C., Goodale, E., Kadis, C., 2016. Do unpaved, low-traffic roads affect bird communities? Acta Oecologica 71, 14–21. https://doi.org/10.1016/j.actao.2016.01.004

McKinney, M.L., 2006. Urbanization as a major cause of biotic homogenization. Biol. Conserv., Urbanization 127, 247–260. https://doi.org/10.1016/j.biocon.2005.09.005

McMahon, B.J., Doyle, S., Gray, A., Kelly, S.B.A., Redpath, S.M., 2020. European bird declines: Do we need to rethink approaches to the management of abundant generalist predators? J. Appl. Ecol. 57, 1885–1890. https://doi.org/10.1111/1365-2664.13695

McWilliams, C., Lurgi, M., Montoya, J.M., Sauve, A., Montoya, D., 2019. The stability of multitrophic communities under habitat loss. Nat. Commun. 10, 2322. https://doi.org/10.1038/s41467-019-10370-2

Meunier, F.D., Verheyden, C., Jouventin, P., 1999. Bird communities of highway verges: Influence of adjacent habitat and roadside management. Acta Oecologica 20, 1–13. https://doi.org/10.1016/S1146-609X(99)80010-1

Morelli, F., Beim, M., Jerzak, L., Jones, D., Tryjanowski, P., 2014. Can roads, railways and related structures have positive effects on birds? – A review. Transp. Res. Part Transp. Environ. 30, 21–31. https://doi.org/10.1016/j.trd.2014.05.006

Morelli, F., Benedetti, Y., Delgado, J.D., 2020. A forecasting map of avian roadkill-risk in Europe: A tool to identify potential hotspots. Biol. Conserv. 249, 108729. https://doi.org/10.1016/j.biocon.2020.108729

Morelli, F., Pruscini, F., Santolini, R., Perna, P., Benedetti, Y., Sisti, D., 2013. Landscape heterogeneity metrics as indicators of bird diversity: Determining the optimal spatial scales in different landscapes. Ecol. Indic. 34, 372–379. https://doi.org/10.1016/j.ecolind.2013.05.021

Muñoz, P.T., Torres, F.P., Megías, A.G., 2015. Effects of roads on insects: a review. Biodivers. Conserv. 24, 659–682. https://doi.org/10.1007/s10531-014-0831-2

Pacifici, K., Simons, T.R., Pollock, K.H., 2008. Effects of vegetation and background noise on the detection process in auditory avian point-count surveys. The Auk 125, 600–607.

Padgham, M., Lovelace, R., Salmon, M., Rudis, B., 2017. osmdata. J. Open Source Softw. 2, 305. https://doi.org/10.21105/joss.00305

Paradis, E., Claude, J., Strimmer, K., 2004. APE: analyses of phylogenetics and evolution in R language. Bioinformatics 20, 289–290.

Phillips, B.B., Bullock, J.M., Osborne, J.L., Gaston, K.J., 2020. Ecosystem service provision by road verges. J. Appl. Ecol. 57, 488–501. https://doi.org/10.1111/1365-2664.13556

Pita, R., Mira, A., Moreira, F., Morgado, R., Beja, P., 2009. Influence of landscape characteristics on carnivore diversity and abundance in Mediterranean farmland. Agric. Ecosyst. Environ. 132, 57–65. https://doi.org/10.1016/j.agee.2009.02.008

Proppe, D.S., Sturdy, C.B., St. Clair, C.C., 2013. Anthropogenic noise decreases urban songbird diversity and may contribute to homogenization. Glob. Change Biol. 19, 1075–1084. https://doi.org/10.1111/gcb.12098

QGIS.org, 2020. QGIS Geographic Information System. Open Source Geospatial Foundation Project. http://qgis.org.

R Development Core Team, 2020. R: A language and environment for statistical computing. R Foundation for Statistical Computing, Vienna, Austria. URL https://www.R-project.org/.

Rosenberg, K.V., Dokter, A.M., Blancher, P.J., Sauer, J.R., Smith, A.C., Smith, P.A., Stanton, J.C., Panjabi, A., Helft, L., Parr, M., Marra, P.P., 2019. Decline of the North American avifauna. Science 366, 120–124. https://doi.org/10.1126/science.aaw1313

Santos, S.M., Mira, A., Salgueiro, P.A., Costa, P., Medinas, D., Beja, P., 2016. Avian trait-mediated vulnerability to road traffic collisions. Biol. Conserv. 200, 122–130. https://doi.org/10.1016/j.biocon.2016.06.004

Schlaepfer, M.A., Runge, M.C., Sherman, P.W., 2002. Ecological and evolutionary traps. Trends Ecol. Evol. 17, 474–480.

Sullivan, B.L., Aycrigg, J.L., Barry, J.H., Bonney, R.E., Bruns, N., Cooper, C.B., Damoulas, T., Dhondt, A.A., Dietterich, T., Farnsworth, A., Fink, D., Fitzpatrick, J.W., Fredericks, T., Gerbracht, J., Gomes, C., Hochachka, W.M., Iliff, M.J., Lagoze, C., La Sorte, F.A., Merrifield, M., Morris, W., Phillips, T.B., Reynolds, M., Rodewald, A.D., Rosenberg, K.V., Trautmann, N.M., Wiggins, A., Winkler, D.W., Wong, W.-K., Wood, C.L., Yu, J., Kelling, S., 2014. The eBird enterprise: An integrated approach to development and application of citizen science. Biol. Conserv. 169, 31–40. https://doi.org/10.1016/j.biocon.2013.11.003

Summers, P.D., Cunnington, G.M., Fahrig, L., 2011. Are the negative effects of roads on breeding birds caused by traffic noise?: Birds and traffic noise. J. Appl. Ecol. 48, 1527–1534. https://doi.org/10.1111/j.1365-2664.2011.02041.x

Trombulak, S.C., Frissell, C.A., 2000. Review of Ecological Effects of Roads on Terrestrial and Aquatic Communities. Conserv. Biol. 14, 18–30. https://doi.org/10.1046/j.1523-1739.2000.99084.x

Tung Ho, L. si, Ané, C., 2014. A Linear-Time Algorithm for Gaussian and Non-Gaussian Trait Evolution Models. Syst. Biol. 63, 397–408. https://doi.org/10.1093/sysbio/syu005

Tylianakis, J.M., Didham, R.K., Bascompte, J., Wardle, D.A., 2008. Global change and species interactions in terrestrial ecosystems. Ecol. Lett. 11, 1351–1363. https://doi.org/10.1111/j.1461-0248.2008.01250.x

Van der Ree, R., Smith, D.J., Grilo, C., 2015. Handbook of road ecology. John Wiley & Sons.

Villemey, A., Jeusset, A., Vargac, M., Bertheau, Y., Coulon, A., Touroult, J., Vanpeene, S., Castagneyrol, B., Jactel, H., Witte, I., Deniaud, N., Flamerie De Lachapelle, F., Jaslier, E., Roy, V., Guinard, E., Le Mitouard, E., Rauel, V., Sordello, R., 2018. Can linear transportation infrastructure verges constitute a habitat and/or a corridor for insects in temperate landscapes? A systematic review. Environ. Evid. 7, 5. https://doi.org/10.1186/s13750-018-0117-3

Whelan, C.J., Wenny, D.G., Marquis, R.J., 2008. Ecosystem services provided by birds. Ann. N. Y. Acad. Sci. 1134, 25–60.

Wood, C., Sullivan, B., Iliff, M., Fink, D., Kelling, S., 2011. eBird: Engaging Birders in Science and Conservation. PLOS Biol. 9, e1001220. https://doi.org/10.1371/journal.pbio.1001220

