## Supplementary material for "Road encroachment mediates species occupancy, trait filtering and dissimilarity of songbird communities"

Supporting information

### S1 – Bird information

Table S1.1 – Number of unique point-count sites where each species was recorded. The data of the species in bold (n=58) was used to test if species are differentially affected by roads, by assessing the relationship between the species’ presence/absence and the road density and environmental predictors.

| **Family** | **Species** | **Common name** | **N** |
| --- | --- | --- | --- |
| Acrocephalidae | ***Hippolais polyglotta*** | **Melodious Warbler** | **339** |
|  | ***Acrocephalus arundinaceus*** | **Great Reed Warbler** | **152** |
|  | ***Acrocephalus scirpaceus*** | **Eurasian Reed Warbler** | **115** |
|  | *Acrocephalus schoenobaenus* | Sedge Warbler | 20 |
|  | *Iduna opaca* | Western Olivaceous Warbler | 14 |
|  | *Acrocephalus baeticatus* | African Reed Warbler | 7 |
|  | *Acrocephalus melanopogon* | Moustached Warbler | 5 |
| Aegithalidae | ***Aegithalos caudatus*** | **Long-tailed Tit** | **197** |
| Alaudidae | ***Galerida cristata*** | **Crested Lark** | **1326** |
|  | ***Lullula arborea*** | **Wood Lark** | **606** |
|  | ***Melanocorypha calandra*** | **Calandra Lark** | **449** |
|  | ***Galerida theklae*** | **Thekla's Lark** | **375** |
|  | ***Calandrella brachydactyla*** | **Greater Short-toed Lark** | **327** |
|  | ***Alauda arvensis*** | **Eurasian Skylark** | **326** |
|  | *Alaudala rufescens* | Lesser Short-toed Lark | 25 |
|  | *Chersophilus duponti* | Dupont's Lark | 10 |
| Certhiidae | ***Certhia brachydactyla*** | **Short-toed Treecreeper** | **583** |
| Cisticolidae | ***Cisticola juncidis*** | **Zitting Cisticola** | **1033** |
| Emberizidae | ***Emberiza calandra*** | **Corn Bunting** | **1689** |
|  | ***Emberiza cirlus*** | **Cirl Bunting** | **269** |
|  | ***Emberiza cia*** | **Rock Bunting** | **225** |
|  | *Emberiza hortulana* | Ortolan Bunting | 28 |
|  | *Emberiza schoeniclus* | Reed Bunting | 19 |
|  | *Emberiza citrinella* | Yellowhammer | 17 |
| Fringillidae | ***Serinus serinus*** | **European Serin** | **1418** |
|  | ***Fringilla coelebs*** | **Common Chaffinch** | **1330** |
|  | ***Carduelis carduelis*** | **European Goldfinch** | **1299** |
|  | ***Chloris chloris*** | **European Greenfinch** | **1046** |
|  | ***Linaria cannabina*** | **Eurasian Linnet** | **900** |
|  | ***Coccothraustes coccothraustes*** | **Hawfinch** | **97** |
|  | *Pyrrhula pyrrhula* | Eurasian Bullfinch | 43 |
|  | *Loxia curvirostra* | Red Crossbill | 38 |
|  | *Spinus spinus* | Eurasian Siskin | 19 |
| Fringillidae | *Carduelis citrinella* | Citril Finch | 16 |
| (cont.) | *Fringilla montifringilla* | Brambling | 4 |
|  | *Haemorhous mexicanus* | House Finch | 2 |
| Leiothrichidae | *Leiothrix lutea* | Red-billed Leiothrix | 6 |
| Locustellidae | *Locustella luscinioides* | Savi's Warbler | 11 |
|  | *Locustella naevia* | Common Grasshopper-Warbler | 2 |
| Motacillidae | ***Motacilla alba*** | **White Wagtail** | **535** |
|  | ***Motacilla flava*** | **Western Yellow Wagtail** | **257** |
|  | *Anthus pratensis^*^* | Meadow Pipit | 165 |
|  | ***Motacilla cinerea*** | **Gray Wagtail** | **93** |
|  | ***Anthus campestris*** | **Tawny Pipit** | **75** |
|  | ***Anthus trivialis*** | **Tree Pipit** | **58** |
|  | *Anthus spinoletta* | Water Pipit | 30 |
| Muscicapidae | ***Luscinia megarhynchos*** | **Common Nightingale** | **1050** |
|  | ***Erithacus rubecula*** | **European Robin** | **818** |
|  | ***Saxicola rubicola*** | **European Stonechat** | **811** |
|  | ***Phoenicurus ochruros*** | **Black Redstart** | **419** |
|  | ***Oenanthe oenanthe*** | **Northern Wheatear** | **125** |
|  | ***Monticola solitarius*** | **Blue Rock-Thrush** | **70** |
|  | *Muscicapa striata* | Spotted Flycatcher | 42 |
|  | *Phoenicurus phoenicurus* | Common Redstart | 35 |
|  | *Oenanthe leucura* | Black Wheatear | 31 |
|  | *Ficedula hypoleuca* | European Pied Flycatcher | 20 |
|  | *Monticola saxatilis* | Rufous-tailed Rock-Thrush | 13 |
|  | *Saxicola rubetra* | Whinchat | 12 |
|  | *Cercotrichas galactotes* | Rufous-tailed Scrub-Robin | 9 |
|  | *Luscinia svecica* | Bluethroat | 4 |
|  | *Ficedula parva* | Red-breasted Flycatcher | 1 |
| Oriolidae | ***Oriolus oriolus*** | **Eurasian Golden Oriole** | **502** |
| Panuridae | *Panurus biarmicus* | Bearded Reedling | 7 |
| Paridae | ***Parus major*** | **Great Tit** | **1060** |
|  | ***Cyanistes caeruleus*** | **Eurasian Blue Tit** | **911** |
|  | ***Periparus ater*** | **Coal Tit** | **327** |
|  | ***Lophophanes cristatus*** | **Crested Tit** | **219** |
|  | *Poecile palustris* | Marsh Tit | 7 |
| Passeridae | ***Passer domesticus*** | **House Sparrow** | **1517** |
|  | ***Passer hispaniolensis*** | **Spanish Sparrow** | **307** |
|  | ***Petronia petronia*** | **Rock Sparrow** | **221** |
|  | ***Passer montanus*** | **Eurasian Tree Sparrow** | **169** |
|  | *Montifringilla nivalis* | White-winged Snowfinch | 2 |
| Phylloscopidae | ***Phylloscopus bonelli*** | **Western Bonelli's Warbler** | **252** |
| Phylloscopidae | ***Phylloscopus ibericus*** | **Iberian Chiffchaff** | **175** |
| (cont.) | ***Phylloscopus collybita*** | **Common Chiffchaff** | **174** |
|  | *Phylloscopus trochilus* | Willow Warbler | 36 |
|  | *Phylloscopus sibilatrix* | Wood Warbler | 3 |
|  | *Phylloscopus humei* | Hume's Warbler | 1 |
| Prunellidae | ***Prunella modularis*** | **Dunnock** | **141** |
|  | *Prunella collaris* | Alpine Accentor | 1 |
| Regulidae | ***Regulus ignicapilla*** | **Common Firecrest** | **238** |
|  | *Regulus regulus* | Goldcrest | 4 |
| Remizidae | *Remiz pendulinus* | Eurasian Penduline-Tit | 32 |
| Scotocercidae | ***Cettia cetti*** | **Cetti's Warbler** | **643** |
| Sittidae | ***Sitta europaea*** | **Eurasian Nuthatch** | **280** |
| Sturnidae | ***Sturnus unicolor*** | **Spotless Starling** | **1502** |
|  | *Sturnus vulgaris* | European Starling | 44 |
|  | *Pastor roseus* | Rosy Starling | 2 |
|  | *Acridotheres cristatellus* | Crested Myna | 1 |
| Sylviidae | ***Sylvia atricapilla*** | **Eurasian Blackcap** | **1025** |
|  | ***Sylvia melanocephala*** | **Sardinian Warbler** | **1011** |
|  | ***Sylvia undata*** | **Dartford Warbler** | **210** |
|  | ***Sylvia cantillans*** | **Subalpine Warbler** | **169** |
|  | ***Sylvia hortensis*** | **Western Orphean Warbler** | **158** |
|  | ***Sylvia communis*** | **Greater Whitethroat** | **115** |
|  | *Sylvia borin* | Garden Warbler | 40 |
|  | *Sylvia conspicillata* | Spectacled Warbler | 30 |
| Tichodromidae | *Tichodroma muraria* | Wallcreeper | 2 |
| Troglodytidae | ***Troglodytes troglodytes*** | **Eurasian Wren** | **879** |
| Turdidae | ***Turdus merula*** | **Eurasian Blackbird** | **1784** |
|  | ***Turdus philomelos*** | **Song Thrush** | **243** |
|  | ***Turdus viscivorus*** | **Mistle Thrush** | **229** |
|  | *Turdus iliacus* | Redwing | 4 |
|  | *Turdus pilaris* | Fieldfare | 2 |
|  | *Turdus torquatus* | Ring Ouzel | 2 |

* *Anthus pratensis* does not breed in Iberia

#### S2 - Species traits

##### S2.1. Trait information

Table S2.1.1 – Species traits (per main group) used to test if roads may act as a filter or evolutionary agent. Trait summary and source are presented.

| **Trait** | **Source** | **Observations** |
| --- | --- | --- |
| ***Physical attributes*** |  |  |
| Body mass | Wilman et al. (2014) | In grams |
| Brain mass | Tsuboi et al. (2018) | Residuals against body mass. |
| ***Environmental tolerance*** |  |  |
| Ground-nesting | Evans et al. (2011) | Species were classified as nesting close to the ground if they typically nested on or just above the ground. |
| City dweller | Maklakov et al. (2011) | Ability to thrive in cities. |
| ***Resource use*** |  |  |
| Invertivory | Wilman et al. (2014) | Percent use of invertebrates in diet. |
| Foraging on the ground | Wilman et al. (2014) | Foraging on the ground is the prevalence (%) of foraging on the ground. |

##### S2.2 - Phylogenetic regression models

The different traits considered are partly correlated due to biogeographic and evolutionary constraints, while the available data for the different traits is different across species. Hence, because the statistical removal of collinearity among multiple traits by multivariate statistical procedures would likely result in a loss of biological interpretability; and filtering the data to obtain all the characteristics for the species collection would imply a drastic reduction in the sample size, we evaluated the relation of road density on each trait separately.

Due to uncertainty in the phylogenetic comparative analysis, we acquired 1,000 equally plausible phylogenetic trees based on the Ericson backbone (Ericson *et al.* 2006) from the BirdTree website (Jetz *et al.* 2012). Using 1000 trees is an adequate number of trees to yield precise parameter estimates (Rubolini *et al.* 2015). We convert each tree to a variance–covariance matrix under the assumption of a Brownian Motion model of character evolution using the ape R package (Paradis *et al.* 2004). Under this assumption, the variance is the branch length from the root to tip and the covariance is the branch length from the root to the most recent common ancestor of each pair of taxa (de Villemereuil *et al.* 2012). Where necessary, we used the *comparative.data* function from the ‘caper’ R package (Orme *et al.* 2018) to subset the phylogenetic trees to the pool of species having information on the respective trait (nesting behaviour and urban tolerance were available for a lower number of species).

##### S2.3 – Species trait data

Table S2.3.1 – Data of species traits used in phylogenetic regression models. Road effect: for each species, the coefficient of the Road variable and its 95% confidence interval (CI) of the respective GLM were used to classify the species with respect to its response to roads as follows: as ‘negatively’ or ‘positively’ associated with road density if the CI was below/above zero, respectively, and as ‘neutral’ if zero was within the CI (see text for details). Body mass in g, diet and foraging in percentage. See table S2.1.1 for trait information and sources.

| **Species** | **Road effect** | **Body mass** | **Brain mass** | **City dweller** | **Ground nester** | **Diet. Inv** | **Forage ground** |
| --- | --- | --- | --- | --- | --- | --- | --- |
| *Acrocephalus arundinaceus* | Negative | 30.0 | 0.87 | No | No | 70 | 20 |
| *Alauda arvensis* | Negative | 37.3 | 0.90 | No | Yes | 40 | 100 |
| *Anthus campestris* | Negative | 23.0 | NA | No | Yes | 70 | 100 |
| *Anthus trivialis* | Negative | 23.3 | 0.59 | No | Yes | 60 | 100 |
| *Calandrella brachydactyla* | Negative | 21.1 | NA | NA | Yes | 60 | 100 |
| *Chloris chloris* | Positive | 26.0 | 0.86 | Yes | No | 0 | 40 |
| *Cisticola juncidis* | Negative | 6.8 | NA | NA | No | 80 | 100 |
| *Cyanistes caeruleus* | Positive | 13.3 | 0.60 | Yes | No | 50 | 10 |
| *Emberiza calandra* | Negative | 48.5 | 1.19 | No | Yes | 30 | 90 |
| *Erithacus rubecula* | Positive | 17.7 | 0.61 | Yes | Yes | 40 | 50 |
| *Fringilla coelebs* | Negative | 23.8 | 0.73 | Yes | No | 60 | 40 |
| *Galerida cristata* | Negative | 42.7 | 1.08 | No | Yes | 40 | 100 |
| *Galerida theklae* | Negative | 38.2 | 0.92 | NA | Yes | 60 | 100 |
| *Hippolais polyglotta* | Negative | 11.0 | NA | NA | No | 80 | 0 |
| *Linaria cannabina* | Negative | 19.5 | 0.66 | Yes | Yes | 20 | 60 |
| *Lophophanes cristatus* | Negative | 11.0 | 0.60 | Yes | No | 60 | 10 |
| *Lullula arborea* | Negative | 26.9 | NA | NA | Yes | 50 | 100 |
| *Melanocorypha calandra* | Negative | 61.6 | NA | No | Yes | 50 | 100 |
| *Motacilla alba* | Positive | 23.9 | 0.56 | Yes | No | 100 | 100 |
| *Motacilla flava* | Negative | 17.7 | 0.43 | No | Yes | 70 | 100 |
| *Oenanthe oenanthe* | Negative | 25.4 | 0.70 | No | Yes | 70 | 100 |
| *Parus major* | Positive | 16.3 | 0.82 | Yes | No | 40 | 0 |
| *Passer domesticus* | Positive | 26.5 | 0.90 | Yes | No | 10 | 50 |
| *Petronia petronia* | Negative | 30.2 | NA | NA | No | 30 | 60 |
| *Phoenicurus ochruros* | Positive | 16.5 | 0.56 | NA | No | 60 | 60 |
| *Prunella modularis* | Negative | 20.2 | 0.67 | Yes | Yes | 50 | 100 |
| *Regulus ignicapilla* | Positive | 5.6 | NA | NA | No | 100 | 0 |
| *Saxicola rubicola* | Negative | 14.1 | 0.47 | No | Yes | 70 | 100 |
| *Serinus serinus* | Positive | 11.2 | 0.57 | Yes | No | 0 | 60 |
| *Sylvia atricapilla* | Positive | 16.7 | 0.60 | Yes | Yes | 50 | 0 |
| *Sylvia cantillans* | Negative | 9.6 | NA | NA | No | 60 | 10 |
| *Sylvia communis* | Negative | 15.1 | 0.50 | No | Yes | 60 | 0 |
| *Sylvia hortensis* | Negative | 21.9 | 0.83 | No | No | 70 | 0 |
| *Sylvia undata* | Negative | 10.8 | NA | NA | No | 70 | 0 |
| *Troglodytes troglodytes* | Positive | 9.7 | 0.48 | Yes | No | 60 | 50 |
| *Turdus merula* | Positive | 102.7 | 1.80 | Yes | No | 50 | 60 |

### References

Ascensão, F., Clevenger, A.P., Grilo, C., Filipe, J. & Santos-Reis, M. (2012). Highway verges as habitat providers for small mammals in agrosilvopastoral environments. *Biodivers. Conserv.*, 21, 3681–3697.

Callaghan, C.T., Major, R.E., Wilshire, J.H., Martin, J.M., Kingsford, R.T. & Cornwell, W.K. (2019). Generalists are the most urban-tolerant of birds: a phylogenetically controlled analysis of ecological and life history traits using a novel continuous measure of bird responses to urbanization. *Oikos*, 0.

Ericson, P.G., Anderson, C.L., Britton, T., Elzanowski, A., Johansson, U.S., Källersjö, M., *et al.* (2006). Diversification of Neoaves: integration of molecular sequence data and fossils. *Biol. Lett.*, 2, 543–547.

Evans, K.L., Chamberlain, D.E., Hatchwell, B.J., Gregory, R.D. & Gaston, K.J. (2011). What makes an urban bird? *Glob. Change Biol.*, 17, 32–44.

Fristoe, T.S., Iwaniuk, A.N. & Botero, C.A. (2017). Big brains stabilize populations and facilitate colonization of variable habitats in birds. *Nat. Ecol. Evol.*, 1, 1706.

Jetz, W., Thomas, G., Joy, J., Hartmann, K. & Mooers, A. (2012). The global diversity of birds in space and time. *Nature*, 491, 444.

Lefebvre, L. & Sol, D. (2008). Brains, Lifestyles and Cognition: Are There General Trends? *Brain. Behav. Evol.*, 72, 135–144.

Lislevand, T., Figuerola, J. & Székely, T. (2007). Avian Body Sizes in Relation to Fecundity, Mating System, Display Behavior, and Resource Sharing. *Ecology*, 88, 1605–1605.

Maklakov, A.A., Immler, S., Gonzalez-Voyer, A., Rönn, J. & Kolm, N. (2011). Brains and the city: big-brained passerine birds succeed in urban environments. *Biol. Lett.*, 7, 730–732.

McKinney, M.L. (2006). Urbanization as a major cause of biotic homogenization. *Biol. Conserv.*, Urbanization, 127, 247–260.

Muñoz, P.T., Torres, F.P. & Megías, A.G. (2015). Effects of roads on insects: a review. *Biodivers. Conserv.*, 24, 659–682.

Orme, D., Freckleton, R., Thomas, G., Petzoldt, T., Fritz, S., Isaac, N., *et al.* (2018). *caper: Comparative Analyses of Phylogenetics and Evolution in R. R package version 1.0.1. https://CRAN.R-project.org/package=caper*.

Paradis, E., Claude, J. & Strimmer, K. (2004). APE: analyses of phylogenetics and evolution in R language. *Bioinformatics*, 20, 289–290.

Pita, R., Mira, A., Moreira, F., Morgado, R. & Beja, P. (2009). Influence of landscape characteristics on carnivore diversity and abundance in Mediterranean farmland. *Agric. Ecosyst. Environ.*, 132, 57–65.

Planillo, A., Kramer-Schadt, S. & Malo, J.E. (2015). Transport Infrastructure Shapes Foraging Habitat in a Raptor Community. *PLOS ONE*, 10, e0118604.

Rubolini, D., Liker, A., Garamszegi, L.Z., Møller, A.P. & Saino, N. (2015). Using the BirdTree. org website to obtain robust phylogenies for avian comparative studies: a primer. *Curr. Zool.*, 61, 959–965.

Santos, S.M., Mira, A., Salgueiro, P.A., Costa, P., Medinas, D. & Beja, P. (2016). Avian trait-mediated vulnerability to road traffic collisions. *Biol. Conserv.*, 200, 122–130.

Tsuboi, M., van der Bijl, W., Kopperud, B.T., Erritzøe, J., Voje, K.L., Kotrschal, A., *et al.* (2018). Breakdown of brain–body allometry and the encephalization of birds and mammals. *Nat. Ecol. Evol.*, 2, 1492–1500.

de Villemereuil, P., Wells, J.A., Edwards, R.D. & Blomberg, S.P. (2012). Bayesian models for comparative analysis integrating phylogenetic uncertainty. *BMC Evol. Biol.*, 12, 102.

Villemey, A., Jeusset, A., Vargac, M., Bertheau, Y., Coulon, A., Touroult, J., *et al.* (2018). Can linear transportation infrastructure verges constitute a habitat and/or a corridor for insects in temperate landscapes? A systematic review. *Environ. Evid.*, 7, 5.

Wilman, H., Belmaker, J., Simpson, J., Rosa, C. de la, Rivadeneira, M.M. & Jetz, W. (2014). EltonTraits 1.0: Species-level foraging attributes of the world’s birds and mammals. *Ecology*, 95, 2027–2027.
